# Accumulation of Oncogenic Mutations During Progression from Healthy Tissue to Cancer

**DOI:** 10.1101/2024.02.18.580841

**Authors:** Ruibo Zhang, Ivana Bozic

## Abstract

Cancers typically result from sequential accumulation of driver mutations in a previously healthy cell. Some of these mutations, such as inactivation of the first copy of a tumor suppressor gene, can be neutral, and some, like those resulting in activation of oncogenes, may provide cells with a selective growth advantage. We study a multi-type branching process that starts with healthy tissue in homeostasis and models accumulation of neutral and advantageous mutations on the way to cancer. We provide results regarding the sizes of premalignant populations and the waiting times to the first cell with a particular combination of mutations, including the waiting time to malignancy. Finally, we apply our results to two specific biological settings: initiation of colorectal cancer and age incidence of chronic myeloid leukemia. Our model allows for any order of neutral and advantageous mutations and can be applied to other evolutionary settings.

## 1 Introduction

Cancer is a genetic disease that results from accumulation of driver mutations which confer a selective growth advantage to tumor cells (Vogelstein and Kinzler 2004). For solid cancers, typically more than one driver mutation is required for the development of malignancy, while a single genetic alteration may be sufficient to cause certain types of leukemia (Vogelstein et al. 2013). With the emergence of advanced sequencing technology, specific driver genes, including ocogenes, tumor suppressor genes and DNA repair genes, have been found to be responsible for carcinogenesis. For example, tumor suppressor genes *APC, TP53* and oncogene *KRAS* are are the most commonly mutated driver genes in colorectal cancer (Fearon 2011; Morin et al. 1997; Tomasetti et al. 2015), and fusion gene *BCR-ABL* is found to cause chronic myeloid leukemia (Deininger et al. 2000).

Some of the key questions in cancer research involve uncovering the identities, the number, the order and the effects of specific driver mutations on tumorigenesis. To facilitate mathematical quantification of the carcinogenic process, stochastic models can be used to model the accumulation of driver mutations, in particular population sizes and arrival time distributions for premalignant and malignant subpopulations. This approach goes back to the multi-stage theory of Armitage and Doll (1954), in which the shape of a cancer age incidence curve is shown to be associated with the required number of driver mutations. More recently, branching processes have been employed to investigate the age incidence of cancer (Meza et al. 2008; Paterson et al. 2020; Wang et al. 2022), cancer relapse and treatment response (Avanzini and Antal 2019; Bozic et al. 2013; Foo et al. 2014; Komarova and Wodarz 2005), and cancer heterogeneity (Durrett et al. 2011).

In the context of cancer initiation, the onset of the process occurs in healthy tissue, when a previously healthy cell receives the first oncogenic alteration. The process proceeds through abnormal growth of the altered subpopulation, acquisition of subsequent driver mutations and further waves of clonal expansion. Previous works that studied accumulation of driver mutations on the way to cancer focused on modeling evolution in exponentially growing populations (Bozic et al. 2010; Durrett and Moseley 2010; Nicholson et al. 2023). These works analyze a process that starts with a single cell that already has selective growth advantage, and model the evolution arising from this single activated cell.

In this paper, we study a process in which the large initial cell population is in homeostasis, capturing the population dynamics both before and during the exponential growth stage. In our model, any sequence of neutral or advantageous genetic alteration can occur and eventually lead to malignancy. Building upon Durrett and Moseley (2010) and Nicholson et al. (2023), we give explicit formulas for population size and arrival time distributions given the order, mutation rates and fitness increments of the driver genes along a specific mutational pathway. Our results are applicable to other multi-hit models that involve the evolution of an initially non-growing population.

## 2 Model

Inspiration for our model comes from initiation of colorectal cancer, which is thought to require inactivation of two tumor suppressor genes and activation of one oncogene (Paterson et al. 2020; Tomasetti et al. 2015; Vogelstein et al. 2013). Tumor suppressor genes, such as *APC* and *TP53*, are the most commonly mutated genes in colorectal cancer, and require inactivation (through genetic alterations) of both alleles to act as cancer driver genes. Oncogenes, such as *KRAS* or *BRAF*, which are also commonly mutated in colorectal cancer, require a single activating mutation in one allele of the gene in question. In other words, initiation of colorectal cancer requires five genetic alterations (two each in two tumor suppressor genes and one in an oncogene). If the first of the five alterations is activation of an oncogene, the crypts carrying that mutation can already exhibit selective growth advantage compared to neighboring crypts, as their rate of crypt fission (division) is significantly increased (Snippert et al. 2014). However, if the first alterations are in tumor suppressor genes, the first one to three alterations may not immediately lead to selective advantage (Paterson et al. 2020). This is because inactivation of a single allele of a tumor suppressor gene typically does not provide selective growth advantage to crypts. Furthermore, some driver genes, such as *TP53*, do not provide selective growth advantage when they are the initial driver alteration, but may lead to abnormal growth if another mutation is subsequently obtained (Paterson et al. 2020).

We study a multi-type branching process generalization of the process above, that starts with a large wild-type population at homeostasis, corresponding to healthy tissue (type 0). As colorectal crypts in homeostasis rarely divide or die (Nicholson et al. 2018), we set the division and death rates of the initial population to 0. In the model, we allow for a number of further oncogenic alterations that initially do not provide selective growth advantage, which occur with distinct constant rates per crypt. After a sufficient number of neutral alterations, the next alteration leads to selective growth advantage in the form of increased division rate. This corresponds, for example, to the inactivation of the second allele of tumor suppressor gene *APC*. After that, subsequent oncogenic mutations, which may be initially neutral, or provide additional selective growth advantage, can accrue. Once a sufficient number of mutations is collected, the crypt becomes malignant. The model can be summarized by the following diagram:

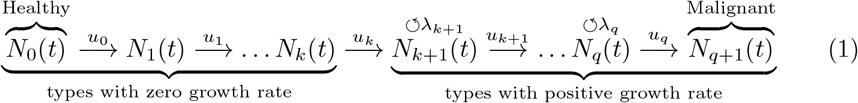

More formally, we study a continuous-time branching process with *q* + 2 types forming a linear evolutionary pathway from type 0 (the healthy type) to type *q* + 1 (the malignant type). We denote the population size of type *i* at time *t* by *N*_*i*_(*t*). The process is started at time 0 with a large healthy population, *N*_0_(0) = *N*. In general, population sizes of individual types may change due to three events: division, death, and mutation. Type *i* cells (or crypts) divide into two daughter cells (crypts) of the same type at rate *b*_*i*_, die at rate *d*_*i*_, and mutate into type *i* + 1 cells at rate *u*_*i*_. We define *λ*_*i*_ := *b*_*i*_ *− d*_*i*_ to be the net growth rate of type *i*.

In the model, the initial type is at homeostasis, with net growth rate *λ*_0_ = *b*_0_ *−d*_0_ = 0. We also assume that the first *k >* 0 mutations that accumulate in the process are neutral, leading to no change in division of death rates. The next mutation provides selective growth advantage, leading to positive net growth rate of the (*k* + 1)-st type, *λ*_*k*+1_ *>* 0. Subsequent mutations may be advantageous or neutral. The main quantity of interest in the model is the waiting time to the first type-*i* cell (crypt)

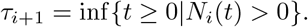

## 3 Results

In this section, we provide analytic results to estimate population sizes and arrival times in the branching process model described in the previous section, and compare them with exact computer simulations of the process. For simplicity, we only discuss the case when mutations are advantageous or neutral and there is no cell death. The scenarios that allow deleterious mutations and cell death are discussed in A.4. We also present two possible applications of the model: initiation of colorectal cancer and incidence of chronic myeloid leukemia.

### 3.1 Population sizes

Individual cells in the model (1) evolve independently. Therefore, the population can be stratified into *N* independent lineages, each of which consists of cells descended from a single original healthy (type 0) cell. Consequently, the population size of a neutral type *l*, 1 *≤ l ≤ k*, counts the number of healthy cells that have evolved to type *l*, but have not changed to type *l* + 1 yet. In particular, at any fixed time *t*, the population of type *l* is distributed as a Binomial(*N, p*(*t*)), with

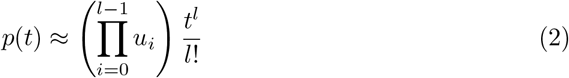

being the time-dependent success probability (for derivation, see A.3). This estimate success probability has a same form as the rate of incidence in Armitage and Doll (1954) with a single unit initial population (see Durrett and Moseley (2010), equation (1)). It follows that, the expectation of *N*_*l*_(*t*) reads

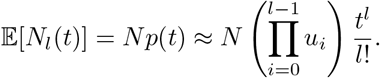

In the small mutation rate regime, *p*(*t*) is a small number, which causes the variance to have a magnitude similar to the mean value:

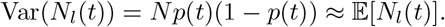

Therefore, the populations of neutral types approximately grow as a power function (Fig. 2).

**Fig. 1.**
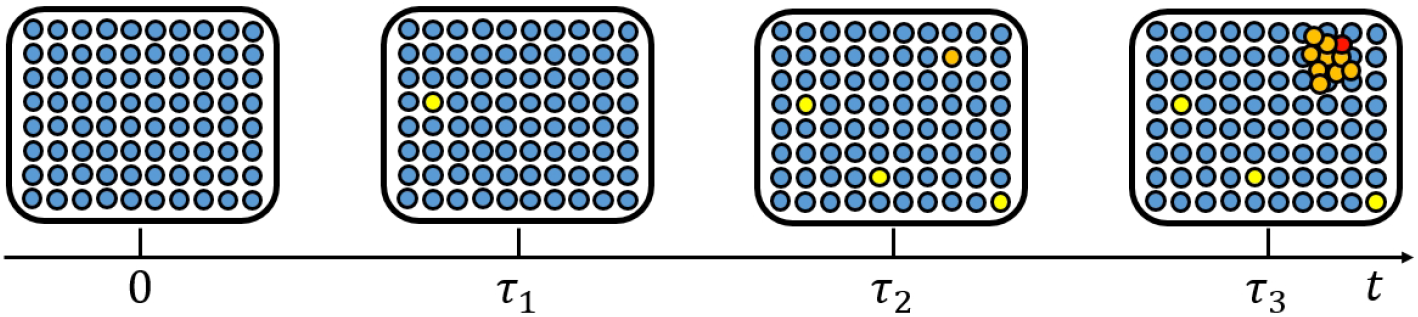
Model illustration. Our model concerns an evolutionary process that starts with a large healthy population (blue circles). In this example, the first oncogenic alteration (yellow) does not provide selective growth advantage. The subsequent genetic alteration (orange) results in growth advantage. Orange cells divide at a higher rate, breaking the homeostasis while still not being considered cancerous. After another genetic alteration takes place, the malignant type (red) emerges.

**Fig. 2.**
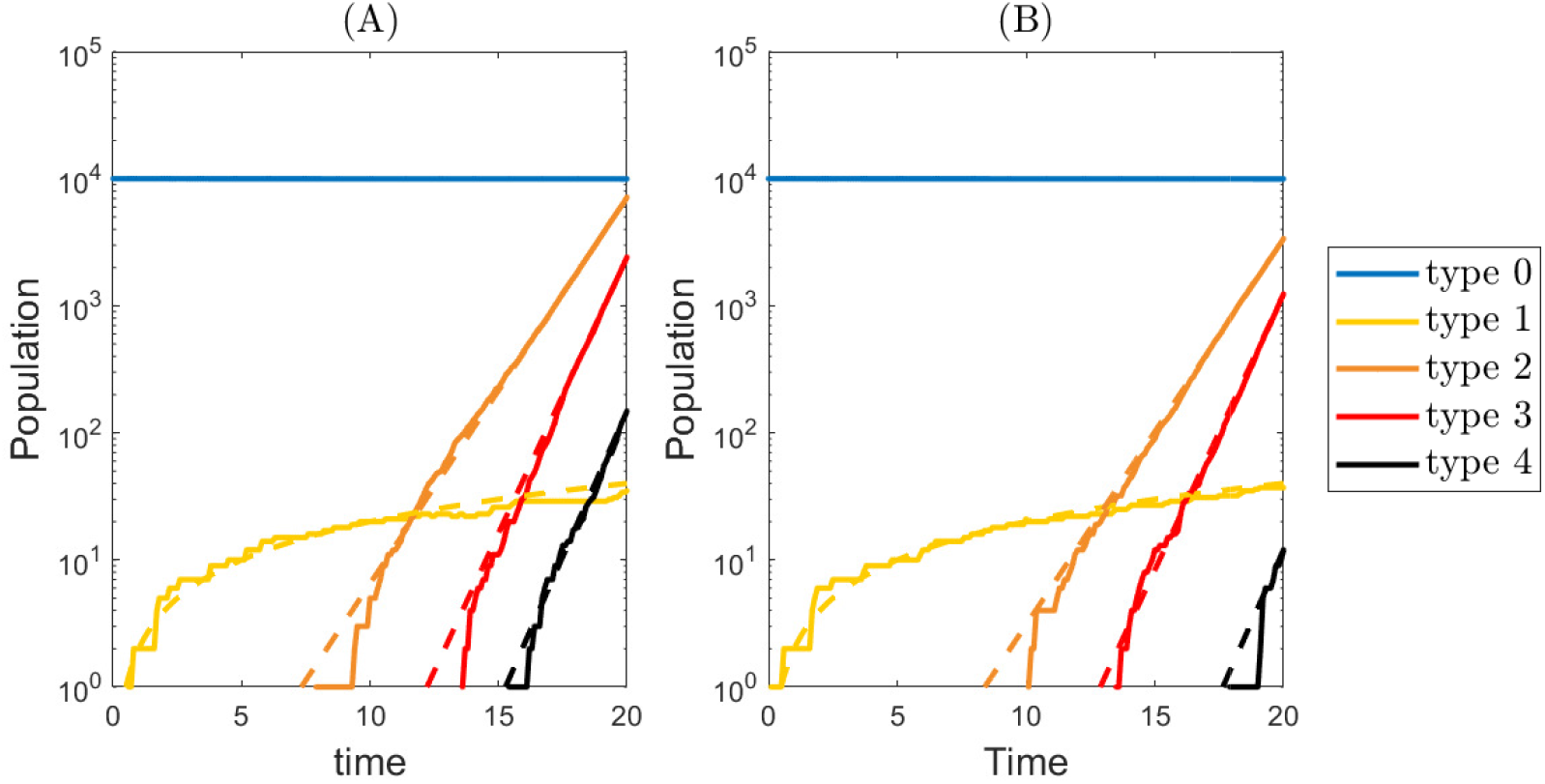
Population sizes in a multi-type branching process. We consider a 5-type branching process with two initial neutral types (with zero growth rate) and three advantageous types (with positive growth rate). Panels (A) and (B) display two different realizations of the process. Solid lines represent computer simulations, and dashed lines represent asymptotic behaviors. Type 1 (light yellow) population grows linearly; Type 2 (orange), type 3 (red), and type 4 (black) populations grow exponentially at large times. Parameter values: *u*_0_ = 2 *×* 10^−4^, *u*_1_ = 8 *×* 10^−3^, *u*_2_ = 5 *×* 10^−3^, *u*_3_ = 6 *×* 10^−3^, *λ*_2_ = 0.7, *λ*_3_ = 1.0, *λ*_4_ = 1.0, *N* = 10^4^, *t* ∈ [0, 20].

Following Durrett and Moseley (2010); Nicholson and Antal (2019); Nicholson et al. (2023), we approximate the population sizes of advantageous types (i.e. types with positive growth rate) in a parameter regime of large times and small mutation rates. For *i ≥* 1, we have shown that

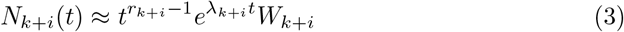

where *r*_*k*+*i*_ = #{*j* = 1, 2, …, |*i*|*λ*_*k*+*j*_ = *λ*_*k*+*i*_} is a constant, and *W*_*k*+*i*_ is a random variable. This approximation separates the stochasity and the time dependence: The population can be decomposed into a multiplication of a time-dependent deterministic function 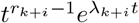 and a time-independent random variable *W*_*k*+*i*_.

Random variable *W*_*k*+*i*_ can be characterized using its Laplace transform:

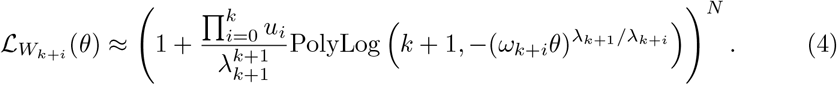

Here *ω*_*k*+*i*_ can be computed iteratively, with *ω*_*k*+1_ = 1, and

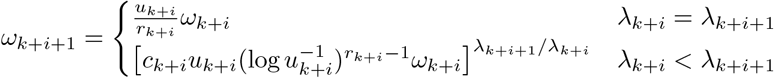

for *i ≥* 1. Finally, we have

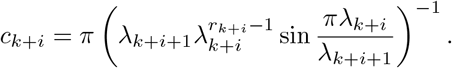

We show that approximations (3) and (4) are in excellent agreement with exact computer simulations of the process in Fig. 2 and Fig. A2.

### 3.2 Arrival times

Before the arrival of the first advantageous type *k* + 1, the total population of the branching process stays fixed. The only possible event for any cell is to change its type into the subsequent type. For a single cell, each alteration requires an exponential waiting time. In a population of cells, the waiting time for a specific type is the minimum time for individual cells to reach that type. This results in the following waiting time distribution for type 1 *≤ l ≤ k* + 1:

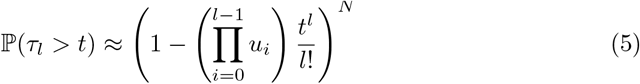

The arrival time of a type that appears after the homeostasis has been partially broken (i.e. whose growth rate is positive) can be split into two segments: (i) The time from the beginning of the process to the arrival of the first advantageous cell, and (ii) The time from the first advantageous cell to the first target type cell. Adapting results from Nicholson et al. (2023), we find an estimate of (ii). Then, treating (i) as a time delay of (ii), we make use of the hypo-exponential distribution to obtain the following approximate formula for the waiting time distribution for type *k* + *i* + 1, *i ≥* 1:

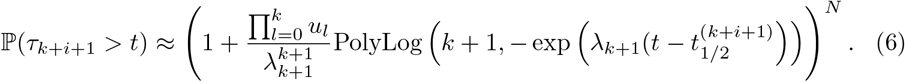

The shape of the waiting time curve is largely determined by *λ*_*k*+1_, the growth rate of the first advantageous type.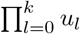 characterizes the amount of time delayed in (i). 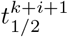 represents the median evolution time from a single type *k* + 1 to the first type *k* + *i* + 1. The value of 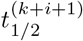 can be derived by the following iterative scheme, which depends on whether (*k* + *i*)-th alteration is neutral or advantageous:

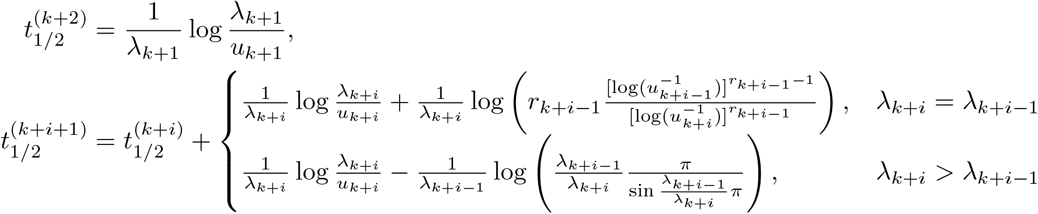

For derivation of results (5) and (6), see A.3. We show that approximations (5) and (6) are in good agreement with exact computer simulations of the process in Fig. 3.

**Fig. 3.**
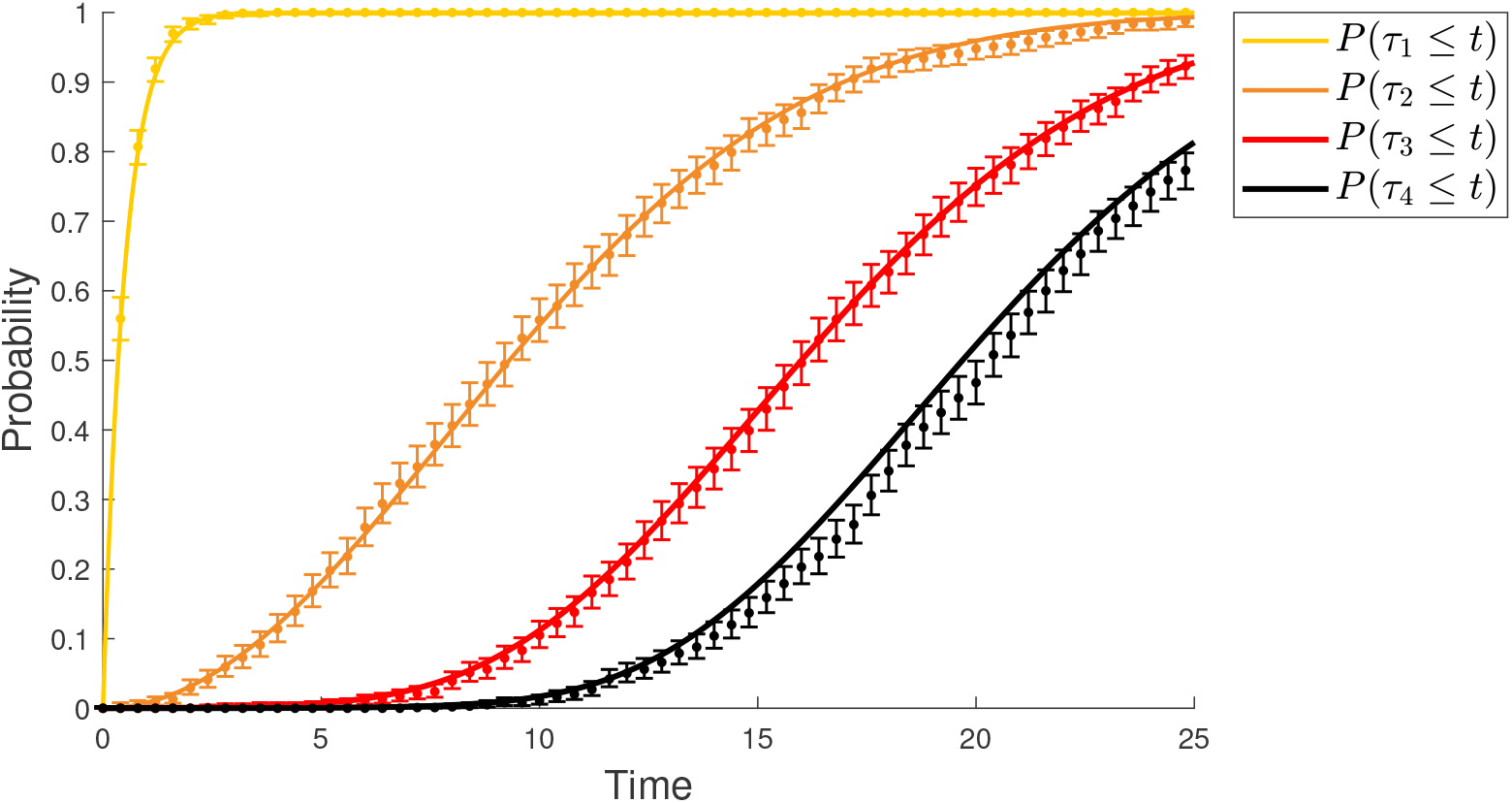
Comparison of analytic results and computer simulations for the waiting time distribution of neutral and advantageous types. Solid lines depict cumulative distribution functions for waiting times to types 1 through 4 in the model described in Fig. 2. Points denote probabilities obtained from computer simulations of the process, with bars showing the 95% confidence interval. Yellow line and orange line show approximation (5); Red line and bleck line show approximation (6). Parameter values: *u*_0_ = 2 *×* 10^−4^, *u*_1_ = 8 *×* 10^−3^, *u*_2_ = 5 *×* 10^−3^, *u*_3_ = 6 *×* 10^−3^, *λ*_2_ = 0.7, *λ*_3_ = 1.0, *λ*_4_ = 1.0, *N* = 10^4^, *t* ∈ [0, 20]. Number of realizations in computer simulation: 1000.

### 3.3 Application: colorectal cancer initiation

Colorectal cancer (CRC) is the end result of a process in which healthy tissue accumulates sequential oncogenic alterations. Multiple driver genes are identified to contribute to this cancerous transformation, but the effect of mutational order of the driver genes on cancer initiation time is not fully understood. Recent work (Paterson et al. 2020) developed a multi-type branching process model to study CRC initiation through acquisition of three common driver genes, tumor suppressors *APC* and *TP53*, and the *KRAS* oncogene. Both alleles of a tumor suppressor gene need to be inactivated for it to function as a driver gene, while only one mutant allele is sufficient for the activation of an oncogene. It follows that CRC initiation involves five sequential genetic alterations. In the model, these genetic alterations may take place through either loss of heterozygosity (LOH) or mutation in any order and at constant rates (Table 1). Zhang et al. (2023) recently studied the waiting time distributions along a single mutational pathway in the order of *APC* inactivation, *KRAS* activation, and *TP53* inactivation.

**Table 1.** CRC driver genes and corresponding parameter values.

| Gene | APC inactivation |  | KRAS activation | TP53 inactivation |  |
| --- | --- | --- | --- | --- | --- |
|  | LOH | Mutation | Mutation | LOH | Mutation |
| rate (per year) | $2.86 \times 10^{-4}$ | $1.06 \times 10^{-5}$ | $9.00 \times 10^{-7}$ | $1.36 \times 10^{-4}$ | $4.56 \times 10^{-7}$ |
| fitness advantage<br>(per year) | 0.20 |  | 0.07 | 0 |  |

Along each individual pathway, population sizes and waiting time distributions can be estimated using the formulas derived in this paper. To demonstrate this, we select two different mutational pathways to CRC and compare our waiting time approximations and the exact computer simulation of the process (Fig. 4). In the first pathway, wild type colonic crypts undergo *APC* inactivation, *KRAS* activation, and *TP53* inactivation consecutively (Fig. 4 (A)). In the second pathway, *APC* inactivation is followed by *TP53* inactivation, and *KRAS* activation (Fig. 4 (B)).

**Fig. 4.**
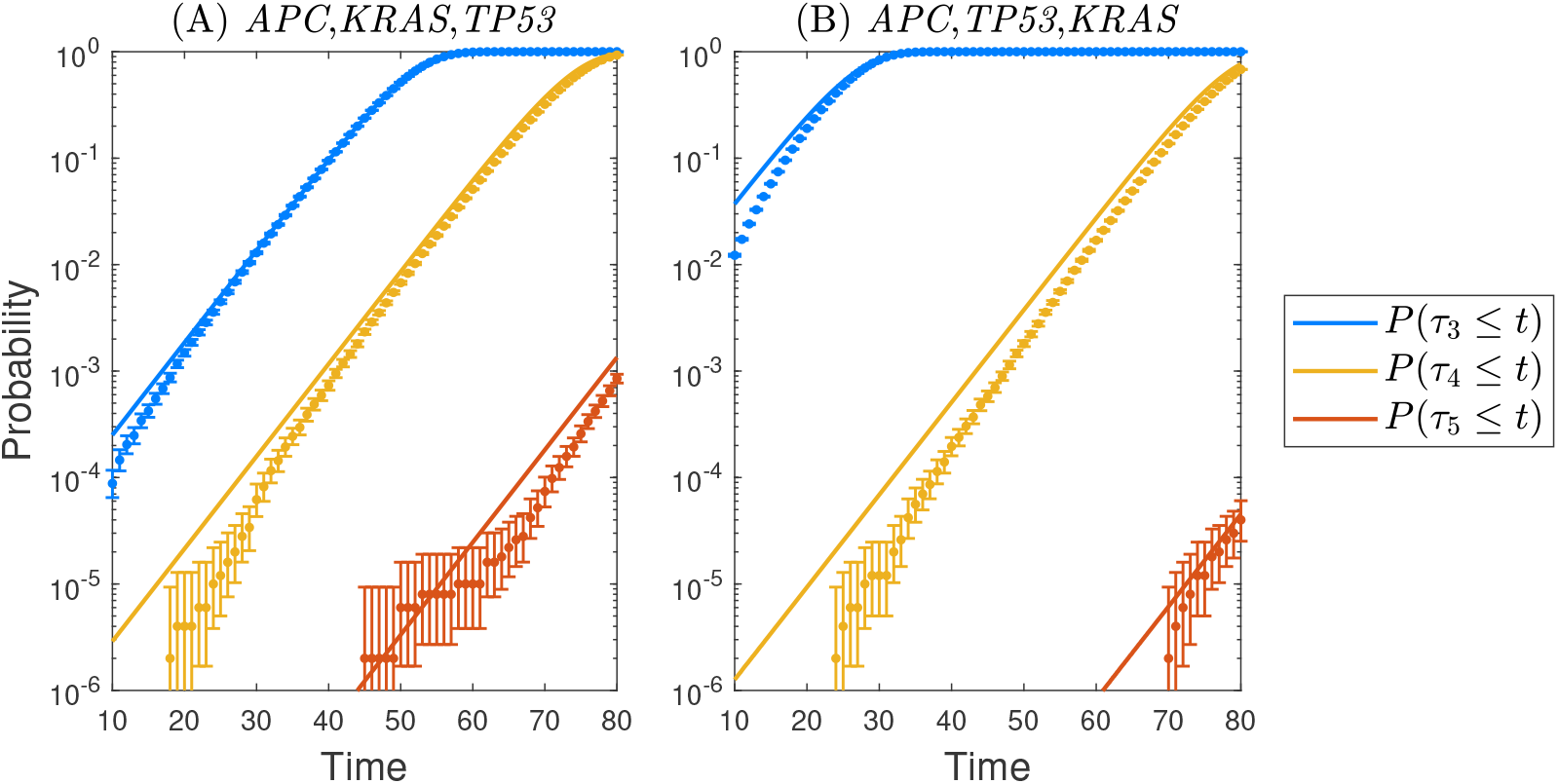
CRC waiting times. Comparison of analytic results and computer simulations for the waiting time distributions of types 3, 4 and 5. Points denote probabilities obtained from computer simulations of the process, with bars showing the 95% confidence interval. Solid lines depict cumulative distribution functions for waiting times obtained from equation (6). In panel (A), the mutational order is *APC* inactivation, *KRAS* activation, and *TP53* inactivation. In panel (B) the mutational order is *APC* inactivation, *TP53* inactivation, and *KRAS* activation. Parameter values: *N* = 10^8^ crypts. Mutation rates and selective growth advantageous are listed in Table 1. Number of realizations in computer simulation: 5 *×* 10^5^.

### 3.4 Application: incidence of chronic myeloid leukemia

Chronic myeloid leukemia (CML) is an uncommon type of cancer that is thought to arise in hematopoietic stem cells. Fusion ocogene *BCR*-*ABL* is identified to initiate the CML carcinogenesis (Deininger et al. 2000). Michor et al. (2006) established a single-hit model that characterizes the malignant transformation of healthy hematopoieticstem cells. In the model, a Moran process is employed to describe the underlining stem cell dynamics. The process starts with a fixed number of healthy stem cells. At each division, a cell is randomly picked and replaced by a newly produced cell, which can carry an oncogenic mutation with some probability. The mutant cell has a selective growth advantage compared to healthy stem cells, leading to clonal expansion of the mutant population. It is assumed that the detection rate of CML is proportional to the population of mutants cells. Michor et al. (2006) derive the detection probability explicitly, fit their model to CML incidence data and conclude that *BCR*-*ABL* alone might be sufficient to initiate CML.

Here, we find that a single-hit branching process model can also recover the CML age incidence curve. To this end, we consider a three-type branching process in which type 0 cells are healthy hematopoietic stem cells, type 1 cells are mutant stem cells with activated BCR-ABL, and type 2 corresponds to CML that has been detected. We assume that healthy stem cells (type 0) are at homeostasis and have a 0 growth rate, and that mutant stem cells (type 1) have a positive growth rate *λ*_1_ *>* 0. In our model, the probability of CML detection at time *t* can be characterized by the type 2 waiting time distribution P(*τ*_2_ *≤ t*). Using equation (6), we find that

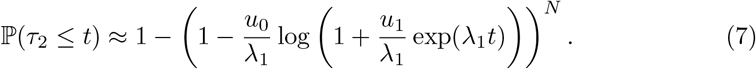

In Fig. 5, we show that equation (7) is in good agreement with the age-prevalence curve of CML.

**Fig. 5.**
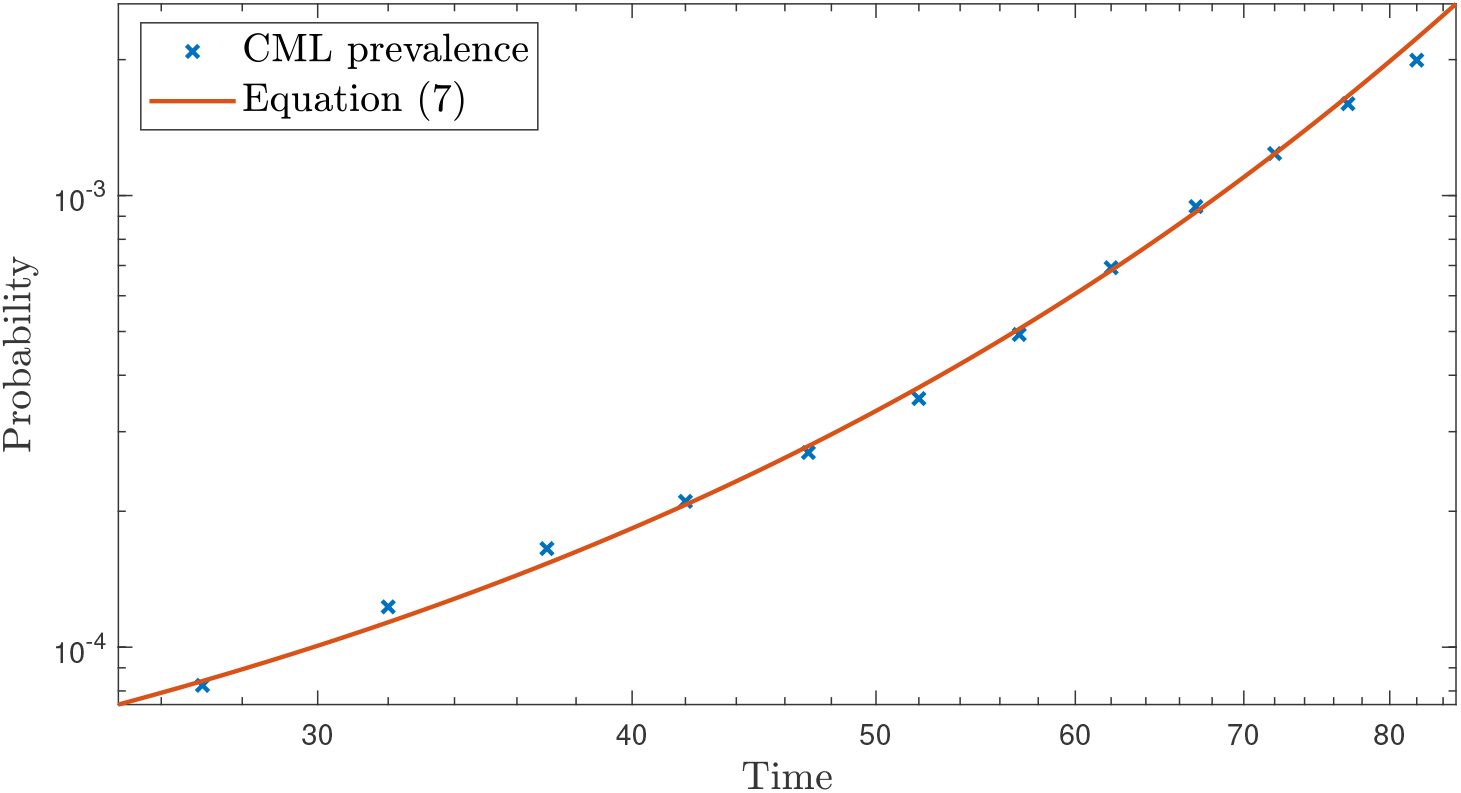
CML incidence. Comparison of the cumulative probability distribution of CML detection (prevalence) from SEER data (Table 1 in Michor et al. (2006)) and equation (7). Parameter values: *N* = 3000, *u*_0_ = 10^−6^, *u*_1_ = 2 *×* 10^−5^, *λ*_1_ = 0.06.

## 4 Discussion

In this work, we study a multi-type branching process that starts with a large cellular population in homeostasis, and models accumulation of neutral and advantageous mutations on the way to malignancy. We derive approximations for population size and arrival time distributions for initial types with no phenotypic changes compared to healthy tissue, as well as for later types that grow abnormally. Applications to modeling the initiation of colorectal cancer and age incidence of chronic myeloid leukemia demonstrate the applicability of our results. Besides cancer evolution, our results are also applicable to other biological phenomena that involve a transformation of a non-growing population through sequential genetic or phenotypic alterations.

We note that the approximations presented here assume that mutation rates are much smaller than growth rates of advantageous types. In particular, for the approximations to be valid, the initial mutation rates have to be small enough compared with the first positive growth rate so that the subsequent mutation occurs when the population of the first advantageous type grows exponentially. This assumption is most likely to be violated when there is a large influx into the first type with a positive growth rate from the previous type resulting in linear population growth when a subsequent mutation occurs.

## Acknowledgments

This work is supported by the National Science Foundation grant DMS-2045166.

## Declarations

The authors declare that they have no conflict of interest.

## Code availability

For access to Gillespie simulation code, please contact the authors.

## Appendix A Methods

In this section, we provide technical details and derive the results presented in the main text. We start by listing assumptions regarding the parameter values that underlie the mathematical proofs. Part of them could be relaxed and will be discussed in A.4. We assume that genetic alterations occur at distinct rates:

### Assumption 1

All the mutation rates are mutually different, i.e. *u*_i_ ≠ *u*_j_, *∀i* ≠ *j*.

We also assume that the mutation rates of genetic alterations are much lower than the growth rates of advantageous mutants. This results in

### Assumption 2

(Small Mutation Rates) *∀*0 *≤ i ≤ q* and *k* + 1 *≤ j ≤ q, u*_i_ *≪ λ*_j_. The *≪* is in the sense that when taking any *u*_i_ *→* 0, *λ*_j_ for any *j* is unaffected.

As we are mainly concerned with neutral and advantageous mutations, we have

### Assumption 3

For all *k* + 1 *≤ i ≤ q −* 1, *λ*_i_ *≤ λ*_i+1_.

Lastly, for simplicity, we initially assume the death rates are zero, i.e.

### Assumption 4

For all *k* + 1 *≤ i ≤ q, d*_i_ = 0.

The last two assumptions (3, 4) are not necessary, and we will discuss the case when these two assumptions do not hold in A.4.

We build upon the work by Nicholson et al. (2023), which provides long-time approximations for population sizes and waiting times in a branching process model with a surviving supercritical initial type. To state the procedure of developing results in this paper, it is necessary to introduce two sub-processes of the main model. Let **e**_*i*_ *∈* ℝ^*q*+2^ be the vector with the (*i* + 1)th coordinate being 1 and all other coordinates being 0, representing the case when the process initial only consists a single type *i* cell. As a Markovian process, the model (1) {**N**(*t*)}_*t≥*0_ is induced by the initial distribution **N**(0) = *N* **e**_0_, i.e. *N* initial healthy cells. We will consider two sub-processes of the main model: (i) a process that starts with a single healthy cell, i.e. 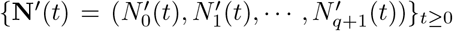 induced by an initial distribution **N**^*′*^(0) = **e**_0_, and (ii) a process that starts with a single type *k* + 1 cell, i.e. 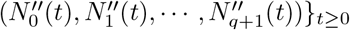 induced by an initial distribution **N**^*′′*^(0) = *e*_*k*+1_.

An outline for obtaining the approximations in this paper includes three steps: First, we employ the results Nicholson et al. (2023) to approximate population sizes and waiting time distributions of *{***N**^*′′*^(*t*)}_*t≥*0_. Next, by using the fact that {**N**^*′*^(*t*)}_*t≥*0_ is essentially a delayed version of {**N**^*′′*^(*t*)}_*t≥*0_, we establish approximations of population sizes and arrival times for {**N**^*′*^(*t*)}_*t≥*0_. Finally, we move from the process with a single initial cell to the model with *N* initial cells and establish our main results by utilizing the branching property.

### A.1 Properties of the model initiated by a single cell with positive growth rate

For {**N**^*′′*^(*t*)*}*_*t≥*0_, approximations for the arrival times and population sizes from type *k* + 1 to type *p* are exhaustively discussed in Nicholson et al. (2023). To apply their results, we introduce the following new notation:

1. *r*_*k*+*i*_ := #{*j* = 1, …, *i* : *λ*_*k*+*k*_ = *λ*_*k*+*i*_*}*: Number of times *λ*_*k*+*i*_ has been attained over types *k* + 1, …, *k* + *i*.
2. 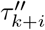 Arrival time until the first type *k* + *i* cell, i.e.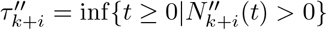
3. 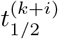 Median arrival time of type *k* + *i* in the process {**N**^*′′*^(*t*)}_*t≥*0_. In other words,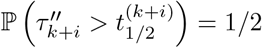.

Under our assumptions (1 - 4), Nicholson et al. (2023) show that there exists (see A.1.1) an approximation of process {**N**^*′′*^(*t*)}_*t≥*0_, denoted by 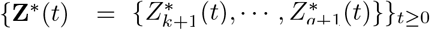 such that 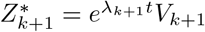 Additionally, for *i ≥* 2,

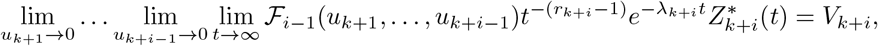

where *ℱ* _*i−*1_ is a known function of the *i−*1 mutations rates and *V*_*k*+*i*_ is a Mittag-Leffler distributed random variable. The Laplace transform of *V*_*k*+*i*_ reads

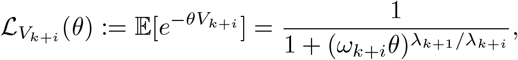

with *ω*_*k*+*i*_ being a constant that depends on the parameters. For computing the value of *ω*_*k*+*i*_, see the recursive formulation after equation (4). The Laplace transforms of *V*_*k*+1_ and *V*_*k*+*i*_ are connected through

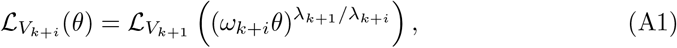

with

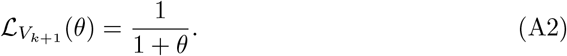

Importantly, the recursive relationship does not depend on the distribution of *V*_*k*+1_ due to the construction of {**Z**^*\**^(*t*)}_*t≥*0_ (see A.1.1). This large-time small-mutation-rate limit leads to the following population size approximation for type *k* + *i*:

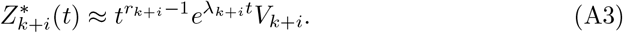

It follows that 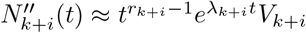 as well.

From this approximation, Nicholson et al. (2023) find that the arrival time of type *k* + *i* + 1, *i ≥* 1 can be estimated by

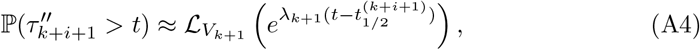

with 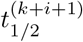 being the median of 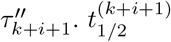 can be expressed using *ω*_*k*+*i*_ by

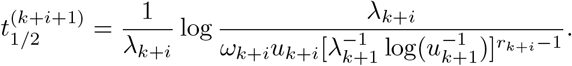

Alternatively, a recursive formulation of 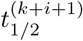 is presented in the main text (see equation (6)).

We note that there is subtle difference between the model in Nicholson et al. (2023) and our model regarding the mutation events. In our model, a mutation from type *i* to type *i* + 1 causes the population of type *i* to decrease by one, i.e. (*i*) ↦ (*i* + 1), while in Nicholson et al. (2023), a mutation events occurs during an asymmetric division in which the type *i* population is not changed, i.e. (*i*) *1→* (*i*)(*i* + 1). Due to the fact that this difference only exists for “growing” types whose net growth rates are assumed to be much greater than mutation rates, Nicholson’s model can be treated as a good approximation for the types *k* + 1 to *q* + 1 in our model.

#### A.1.1 Construction of {Z^*\**^(*t*)}_*t≥*0_

The general strategy of constructing the approximation {**Z**^*\**^(*t*)}_*t≥*0_ was first introduced by Durrett and Moseley (2010) and recently studied by Nicholson et al. (2023). A rigorous mathematical description is presented in Nicholson et al. (2023). Briefly speaking, in the construction, we use a random variable *V*_*i*_ to establish a two-type stochastic process 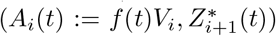, where 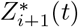 counts a Poisson process with intensity *u*_*i*_*A*_*i*_(*t*) for each single realization of *A*_*i*_(*t*). *V*_*i*_ is found by taking a large time limit of *f* (*t*)^*−*1^*Z*_*i*_(*t*), so that *A*_*i*_(*t*) is a good approximation of *Z*_*i*_(*t*). After that,*V*_*i*+1_ is found to be a large time limiting random variable of 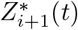, and one con-structs 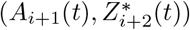 by the same methodology. Nicholson et al. (2023) established results that incorporate small mutation rate limits in the approximations. In our case, the construction starts at type *k* + 1 where 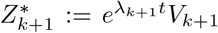 is imposed. Importantly, Nicholson et al. (2023) indicates that the population dynamics of {**Z**^*\**^(*t*)}_*t≥*0_, in a small transition rate limit, is fully induced by the initial type, i.e.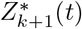. Furthermore, in the approximate model, the probability distribution of the waiting time to type 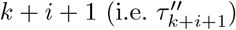 can be expressed by 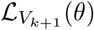 directly (equation (A4)).

### A.2 Properties of the model that starts with a single cell with a zero growth rate

Now we move our focus to the process {**N**^*′*^(*t*)}_*t≥*0_, which represents the branching process that starts with a single type 0 cell. Recall that the growth rates for type 0 through type *k* are zero. This indicates that, the process {**N**^*′*^(*t*)}_*t≥*0_ becomes {**N**^*′′*^(*t*)}_*t≥*0_ after the initial cell collects the first *k* + 1 mutations and changes its type into type *k* + 1. Let 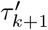 be the arrival time of type *k* + 1 in {**N**^*′*^(*t*)}_*t≥*0_. We find that for *i ≥* 1

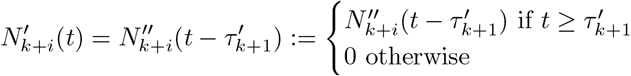

and for 1 *≤ l ≤ k*

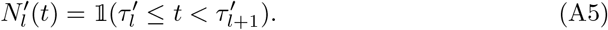

Due to the above expression of 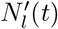, for any 1 *≤ l ≤ k*, the population size follows a Bernoulli distribution with a time-dependent parameter

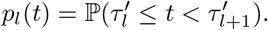

Thus, to understand the properties of **N**^*′*^, we need to obtain the distribution of 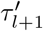, 1 *≤ l ≤ k*. Since each mutation among the first *k* + 1 neutral mutations takes an exponentially distributed time to occur, we have

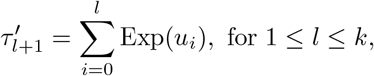

where Exp(*u*_*i*_) denotes an exponentially distributed random variable with density 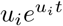. From our assumption 1, *u*_*i*_ ≠ *u*_*j*_ whenever *i* ≠ *j*. Thus,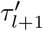follows a hyperexponential distribution with density

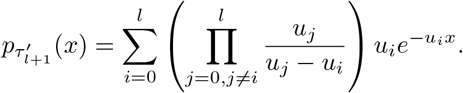

Using this density function, we find that

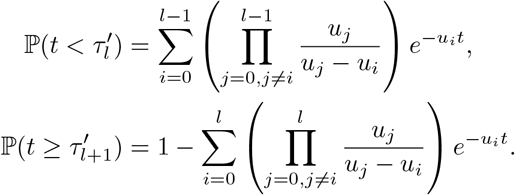

Thus, it follows that the Bernoulli parameter *p*(*t*) for the neutral population is given by

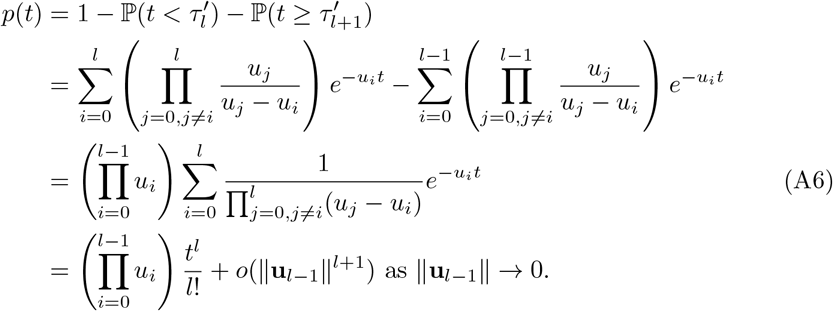

In the last equation, we have used a Taylor expansion in **u**_*l−*1_ := (*u*_0_, *u*_1_, *· · ·, u*_*l−*1_) at 0.

Next, we want to identify the large-time small-mutation-rate population behavior for 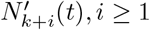, *i ≥* 1. To do that, we first observe that 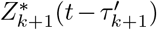 admits a large-time small-mutation-rate limit.

#### Lemma 1

*For the process* 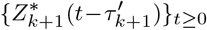, *the following large-time small-mutation-rate limit exists almost surely:*

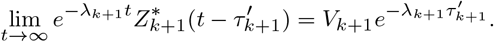

*Proof* Notice that 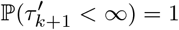. Thus, for each realization *ω*, we have that

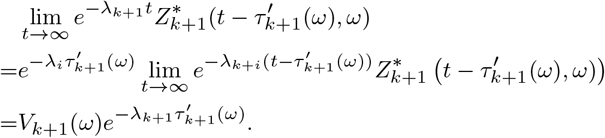

This shows that a large-time small-mutation-rate limit still exists after 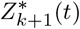 has been shifted by a waiting time 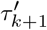. □

We denote the above new limiting random variable by 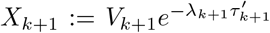 Using 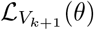 (equation (A2)), we find the Laplace transform of *X*_*k*+1_

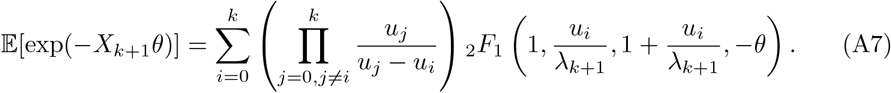

Next, let 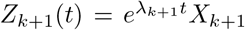 be the approximation of 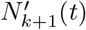. We construct anauxiliary process {**Z**(*t*) = (*Z*_*k*+1_(*t*), *Z*_*k*+2_(*t*), *· · ·, Z*_*k*+*q*_(*t*))}_*t≥*0_ following the procedure described in A.1.1. As an approximation of 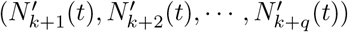, the auxiliary process {**Z**(*t*)}_*t≥*0_ suggests that

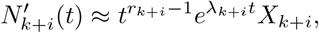

where

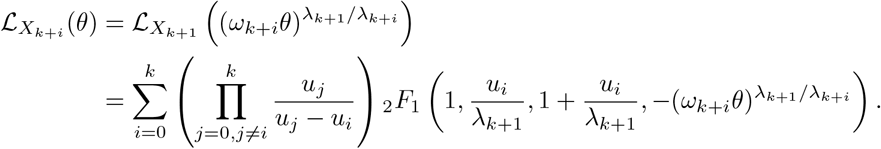

The constant *ω*_*k*+*i*_ is given by the recursive relationship after equation (4). In addition, the {*Z*(*t*)}_*t≥*0_ construction also guarantees that

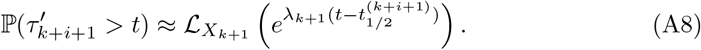

Taking advantage of (A8), we get

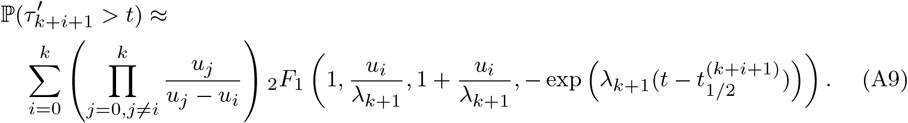

We find that (A9) can be simplified in the small *u*_0_, *· · ·, u*_*k*_ regime, leading to

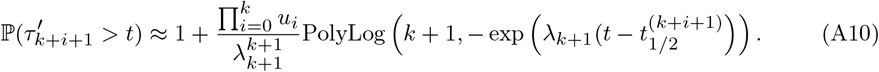

The derivation of equation A10 is provided in Section B.

### A.3 Properties of the model that starts with a large non-growing population

Finally, we consider the population dynamics of {**N**(*t*)}_*t≥*0_ that starts with *N* type 0 cells. By the branching property, {**N**(*t*)}_*t≥*0_ and {**N**^*′*^(*t*)}_*t≥*0_ are related through:

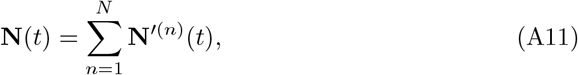

where {{**N**^*′*(1)^(*t*)}_*t≥*0_, *· · ·*, {**N**^*′*(*N*)^(*t*)}_*t≥*0_} is a collection of independent processes that are identically distributed as {**N**^*′*^(*t*)}_*t≥*0_. Equality (A11) reflects that, in a multitype branching process model, each individual in the initial population evolves independently.

For types with zero growth rates, we find that for any 1 *≤ l ≤ k*,

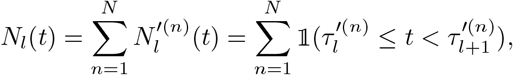

where 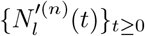 are i.i.d. copies of the process 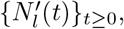 and 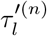 are i.i.d. copies Of 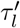. This expression indicates that the population of type *l* (with zero growth rate) at time *t* follows a Binomial (*N, p*(*t*)) distribution. We show that this result is in good agreement with exact computer simulations of the process in Fig. A1.

**Fig. A1.**
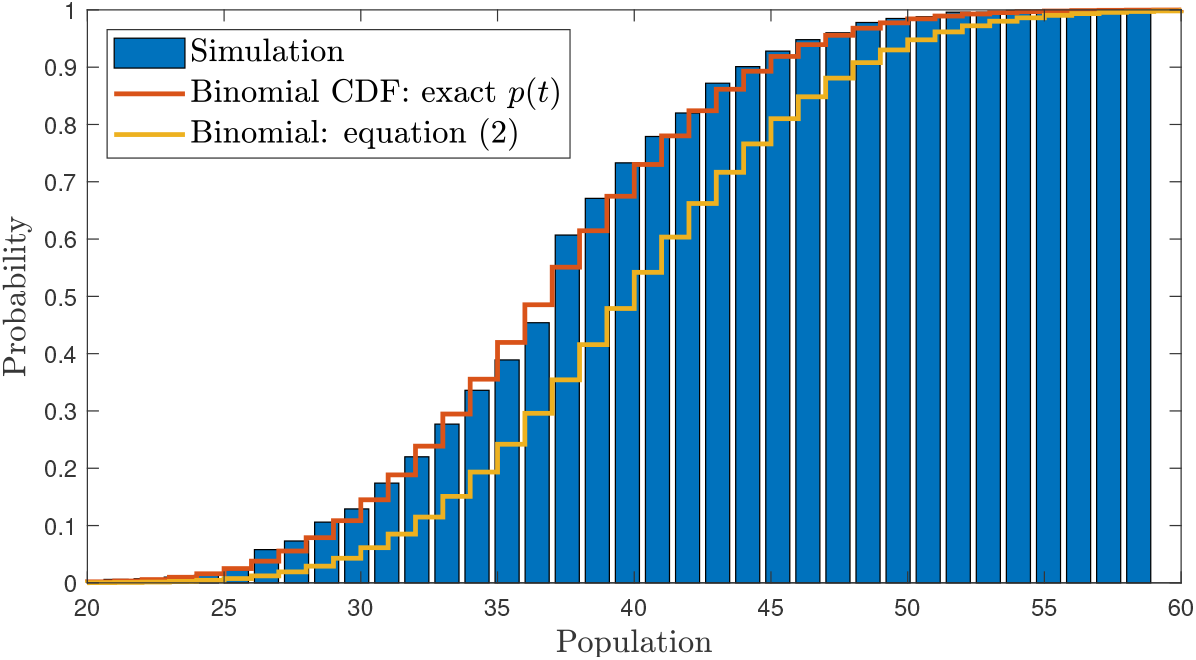
Population dize distribution of a non-growing population. In the model described in Fig. 2, type 1 does not have a selective growth advantage. The simulated cumulative distribution (CDF) of type 1 population at *t* = 20 is presented by the blue bars. Binomial CDFs with exact theoretical success probability *p*(*t*) (A6) and approximate *p*(*t*) (2) are presented in solid lines. Parameter values: *u*_0_ = 2 *×* 10^−4^, *u*_1_ = 8 *×* 10^−3^, *u*_2_ = 5 *×* 10^−3^, *u*_3_ = 6 *×* 10^−3^, *λ*_2_ = 0.7, *λ*_3_ = 1.0, *λ*_4_ = 1.0, *N* = 10^4^, *t* ∈ [0, 20]. Number of realizations in computer simulation: 1000.

The arrival time of type *l* can be treated as the minimum of the type *l* arrival times among the processes that each start with a single cell, i.e.

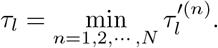

It follows that 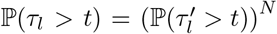. Since 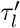 is hypo-exponentially distributed,we find

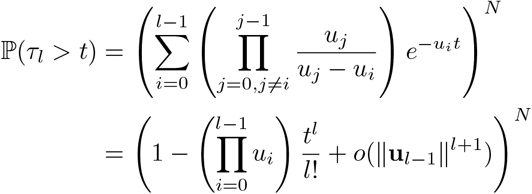

For types with positive growth rates, relationship (A11) indicates that

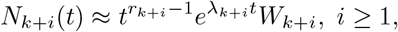

where 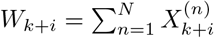 and 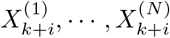 are i.i.d. copies of *X*_*k*+*i*_. The Laplace transform of *W*_*k*+*i*_ is found to be

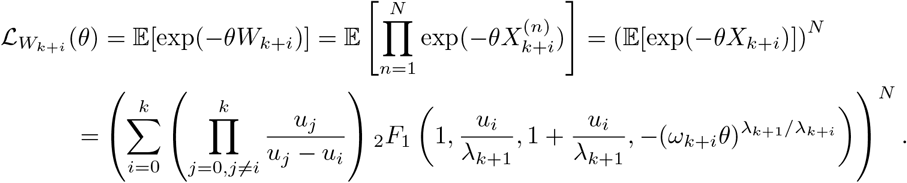

We find an approximate version of 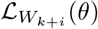 in the small mutation rate paramter regime

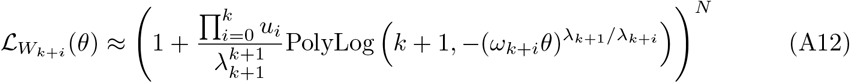

For the arrival time of post-advantageous types, we take advantage of the relationship (A8) and simplification (A10) and to get the distribution of *τ*_*k*+*i*+1_

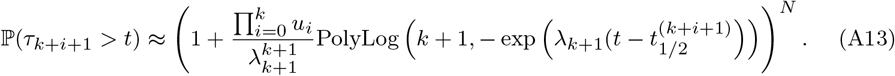

**Fig. A2.**
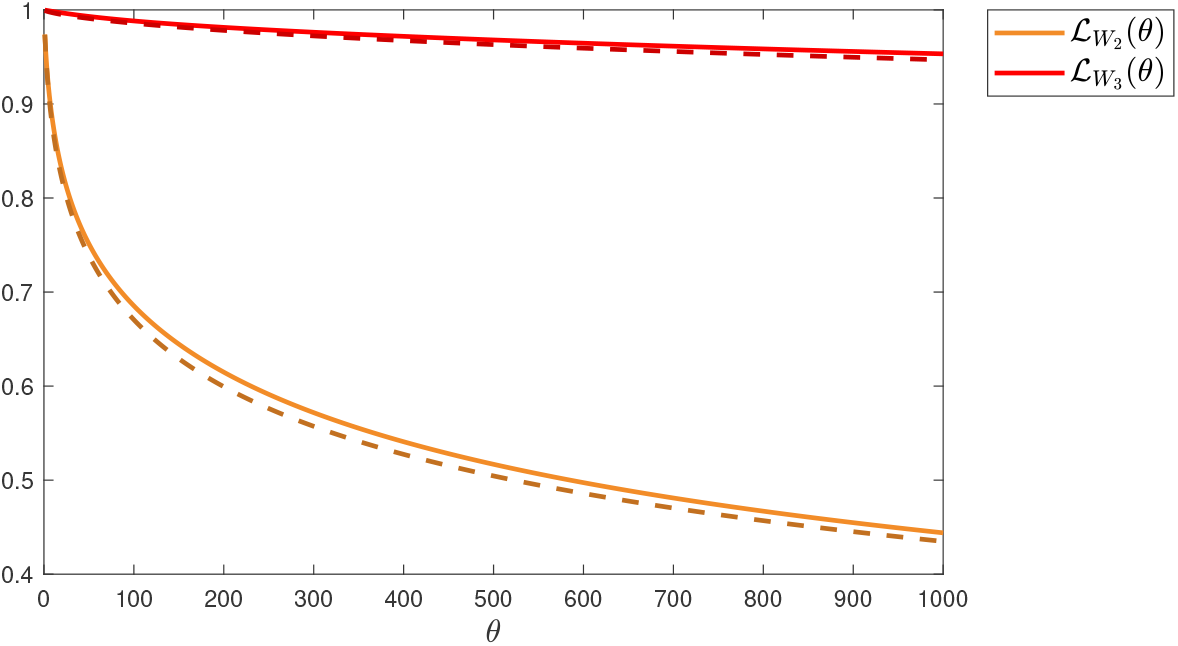
Random amplitude. Laplace transforms of *W*_2_ and *W*_3_ obtained from equation (4) and computer simulations of the process described in Fig. 2. Solid lines depict the simulated Laplace transforms, which are obtained by computing the Laplace transform of scaled populations at *t* = 25. Dashed lines show formula (4). Parameter values: *u*_0_ = 2 *×* 10^−4^, *u*_1_ = 8 *×* 10^−3^, *u*_2_ = 5 *×* 10^−3^, *u*_3_ =6 *×* 10^−3^, *λ*_2_ = 0.7, *λ*_3_ = 1.0, *λ*_4_ = 1.0, *N* = 10^4^, *t* ∈ [0, 20]. Number of realizations in computer simulation: 1000.

### A.4 Allowing death and fitness decreasing events

Following Nicholson et al. (2023), we allow deleterious mutations and positive death rates in the model (1) after the first advantageous mutation as long as the first type with non-zero growth rate is supercritical. Below, we list the results that reflect this parameter regime relaxation. We first introduce the following notation:

1. *δ*_*k*+*i*_ := max_*j*=0,1,*···, i*_{*λ*_*k*+*j*_}: Running-max fitness.
2. *s*_*k*+*i*_ := #{*j* = 1, …, *i* : *λ*_*k*+*j*_ = *λ*_*k*+*i*_}: Number of times *δ*_*i*_ has been attained over types *k* + 1, …, *k* + *i*.

Note that when assumption 3 holds, type *i* always has the largest growth rate among all types before *i*. Thus, *r*_*k*+*i*_ = *s*_*k*+*i*_, *λ*_*k*+1_ = *δ*_*k*+*i*_.

The waiting time distribution of type *k* + *i* + 1 can be approximated in a form similar to equation (6):

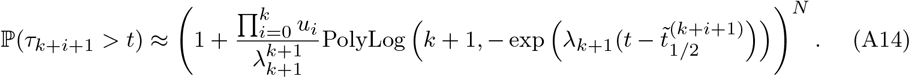

The only difference between (6) and (A14) is that the median time has been changed to 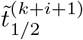. Here, we have (see displays (2) and (5) in Nicholson et al. (2023))

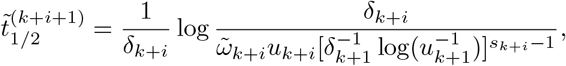

with 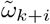 satisfying 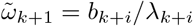 and

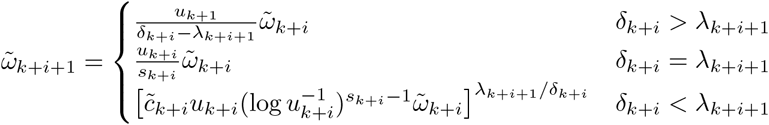

for *i ≥* 1. Finally, we have

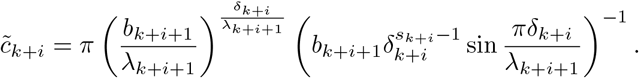

## Appendix B Derivation of the simplified distribution function

In this section, we derive equation (A10), an approximation of equation (A9), using Taylor expansion of hypergeometric functions.

### B.1 The leading order term of the distribution function

The main goal is to find the leading order term of

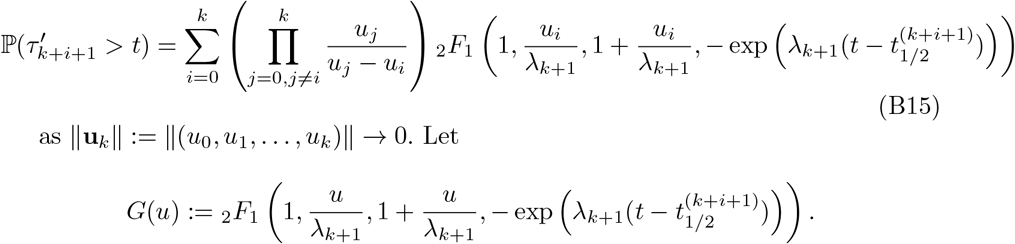

Then (B15) can be written as

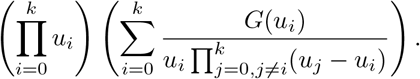

Now, consider the Taylor expansion of *G* at *u* = 0, we have

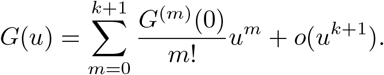

Thus, we have that

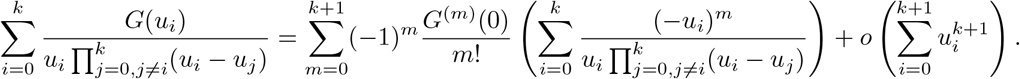

Then, by the fact that (see equation (7) in Supplement File (1) of Bozic et al. (2013))

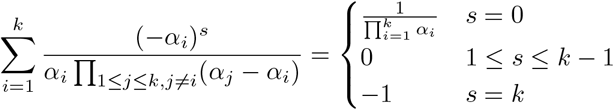

we have

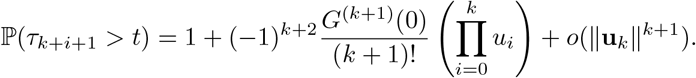

In the next section, we will give an explicit expression of *G*^(*k*+1)^(0).

### B.2 Computing the partial derivatives of the hypergeometric function at a particular point

Here we discuss the derivatives of function *G* at 0. Let

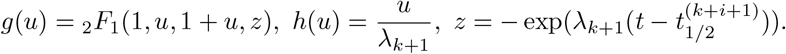

It follows that

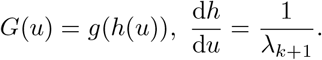

Thus, we have

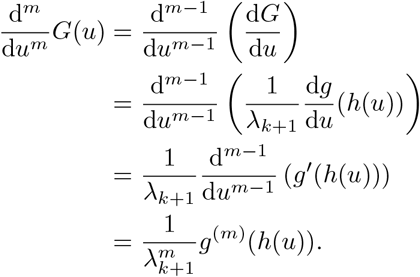

Next, the derivative of *g* is given by the following lemma.

#### Lemma 2

*Let* _2_*F*_1_(*a, b, c, z*) *be the Hypergeometric function and define*

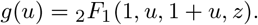

*Then we have*

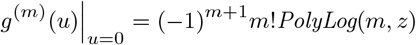

*where PolyLog*(*m, z*) *denotes the PolyLogrithm* (DLMF 2022, 25.12.10).

With this lemma, we immediately see that

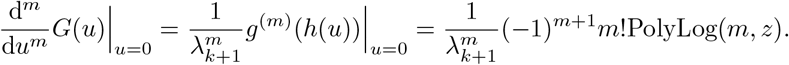

*Proof* Following Ancarani and Gasaneo (2009), to find the *m*th derivative of *g*, we define *F* (*u, z*) := _2_*F*_1_(1, *u*, 1 + *u, z*) and apply the hypergeometric differential equation. The hypergeometric function satisfies the following second-order ODE:

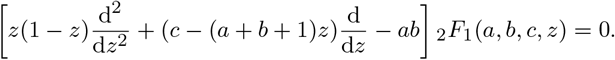

Thus, for *F*, we have that

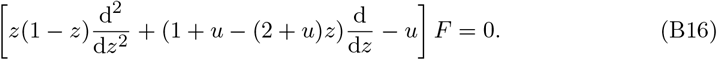

Next, since *F* is analytic in *u*, we can take a derivative with respect to *u* on both sides, which results in

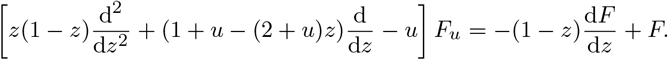

Then, by the formula (DLMF 2022, 15.5.21)

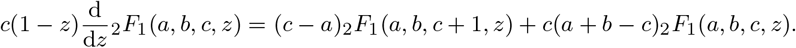

we get that

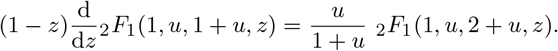

Let 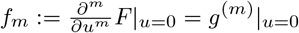. If we look at the derivative at *u* = 0 on both sides, we get

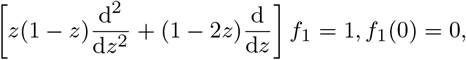

where we have used the fact that _2_*F*_1_(1, 0, 1, *z*) ≡ 1. The above equation is a second order linear equation, but one can see that by letting 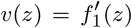, the ODE reduces to a first order equation. Hence, we get a unique solution

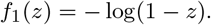

For a general *m*, the ODE for *f*_m_ is

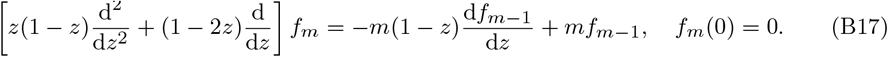

This can be obtained by taking derivatives of the both sides of (B16) *m* times. Let us denote

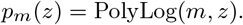

We use induction to show that

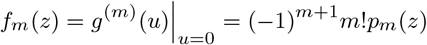

satisfies (B17). For the base case when *m* = 1, we have proved above that

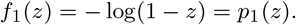

Next, we move to the induction part. Suppose that our claim holds true for *m*, that is

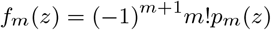

solves (B17). Then, the (*m* + 1)-st equation reads

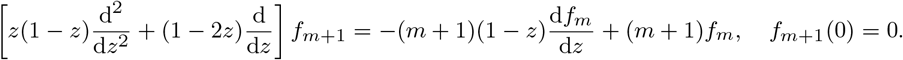

From the derivative formula of the polylogrithm (see (18) at Polylogarithm), we have that

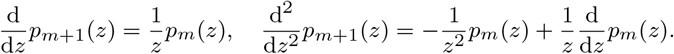

It follows that

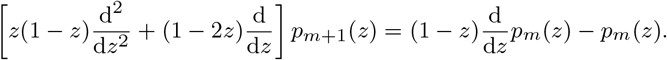

Thus, we see that

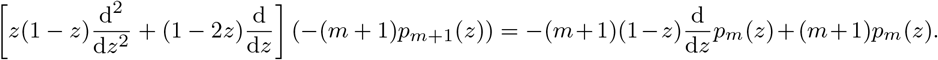

Multiplying both sides by (*−*1)^*m*+1^*m*!, we have

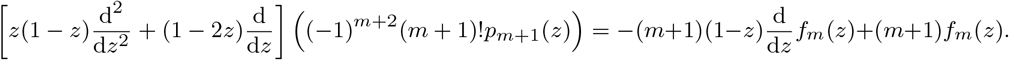

This shows that *f*_m+1_(*z*) = (*−*1)^*m*+2^(*m* + 1)!*p*_m+1_(*z*) and finishes the induction. Finally, we conclude that

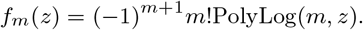

□

## References

Ancarani LU, Gasaneo G (2009) Derivatives of any order of the gaussian hypergeo-metric function 2F1(a, b, c; z) with respect to the parameters a, b and c. J. Phys. A Math. Theor. 42(39):395208. 10.1088/1751-8113/42/39/395208

Armitage P, Doll R (1954) The age distribution of cancer and a multi-stage theory of carcinogenesis. Br. J. Cancer 8(1):1–12. 10.1038/bjc.1954.1

Avanzini S, Antal T (2019) Cancer recurrence times from a branching process model. PLoS Comput. Biol. 15(11):e1007423. 10.1371/journal.pcbi.1007423

Bozic I, Antal T, Ohtsuki H, Carter H, Kim D, Chen S, Karchin R, Kinzler KW, Vogelstein B, Nowak MA (2010) Accumulation of driver and passenger mutations during tumor progression. Proc. Natl. Acad. Sci. U.S.A. 107(43):18545–18550. 10.1073/pnas.1010978107

Bozic I, Reiter JG, Allen B, Antal T, Chatterjee K, Shah P, Moon YS, Yaqubie A, Kelly N, Le DT, Lipson EJ, Chapman PB, Diaz Luis A J, Vogelstein B, Nowak MA (2013) Evolutionary dynamics of cancer in response to targeted combination therapy. eLife 2:e00747. 10.7554/eLife.00747

Deininger MW, Goldman JM, Melo JV (2000) The molecular biology of chronic myeloid leukemia. Blood 96(10):3343–3356. 10.1182/blood.V96.10.3343

DLMF (2022) Nist digital library of mathematical functions. Release 1.1.8 of 2022-12-15, http://dlmf.nist.gov/

Durrett R, Foo J, Leder K, Mayberry J, Michor F (2011) Intratumor heterogeneity in evolutionary models of tumor progression. Genetics 188(2):461–477. 10.1534/genetics.110.125724

Durrett R, Moseley S (2010) Evolution of resistance and progression to disease during clonal expansion of cancer. Theor. Popul. Biol. 77(1):42–48. 10.1016/j.tpb.2009.10.008

Fearon ER (2011) Molecular genetics of colorectal cancer. Annu. Rev. Pathol. 6:479–507. 10.1146/annurev-pathol-011110-130235

Foo J, Leder K, Zhu J (2014) Escape times for branching processes with random mutational fitness effects. Stoch. Process. Their Appl. 124(11):3661–3697. 10.1016/j.spa.2014.06.003

Komarova NL, Wodarz D (2005) Drug resistance in cancer: principles of emergence and prevention. Proc. Natl. Acad. Sci. U.S.A. 102(27):9714–9719. 10.1073/pnas.0501870102

Meza R, Jeon J, Moolgavkar SH, Luebeck EG (2008) Age-specific incidence of cancer: Phases, transitions, and biological implications. Proc. Natl. Acad. Sci. U.S.A. 105(42):16284–16289. 10.1073/pnas.0801151105

Michor F, Iwasa Y, Nowak MA (2006) The age incidence of chronic myeloid leukemia can be explained by a one-mutation model. Proc. Natl. Acad. Sci. U.S.A. 103(40):14931–14934. 10.1073/pnas.0607006103

Morin PJ, Sparks AB, Korinek V, Barker N, Clevers H, Vogelstein B, Kinzler KW (1997) Activation of β-catenin-Tcf signaling in colon cancer by mutations in βcatenin or APC. Science 275(5307):1787–1790. 10.1126/science.275.5307.1787

Nicholson AM, Olpe C, Hoyle A, Thorsen AS, Rus T, M Colomb’se, Brunton-Sim R, Kemp R, Marks K, Quirke P, et al. (2018) Fixation and spread of somatic mutations in adult human colonic epithelium. Cell stem cell 22(6):909–918. 10.1016/j.stem.2018.04.020

Nicholson MD, Antal T (2019) Competing evolutionary paths in growing populations with applications to multidrug resistance. PLoS Comput. Biol. 15:1–25. 10.1371/journal.pcbi.1006866

Nicholson MD, Cheek D, Antal T (2023) Sequential mutations in exponentially growing populations. PLoS Comput. Biol. 19(7):1–32. 10.1371/journal.pcbi.1011289

Paterson C, Clevers H, Bozic I (2020) Mathematical model of colorectal cancer initiation. Proc. Natl. Acad. Sci. U.S.A. 117(34):20681–20688. 10.1073/pnas.2003771117

Snippert HJ, Schepers AG, Van Es JH, Simons BD, Clevers H (2014) Biased competition between Lgr5 intestinal stem cells driven by oncogenic mutation induces clonal expansion. EMBO Rep. 15(1):62–69. 10.1002/embr.201337799

Tomasetti C, Marchionni L, Nowak MA, Parmigiani G, Vogelstein B (2015) Only three driver gene mutations are required for the development of lung and colorectal cancers. Proc. Natl. Acad. Sci. U.S.A. 112(1):118–123. 10.1073/pnas.1421839112

Vogelstein B, Kinzler KW (2004) Cancer genes and the pathways they control. Nat. Med. 10(8):789–799. 10.1038/nm1087

Vogelstein B, Papadopoulos N, Velculescu VE, Zhou S, Diaz Jr LA, Kinzler KW (2013) Cancer genome landscapes. Science 339(6127):1546–1558. 10.1126/science.1235122

Wang Y, Boland CR, Goel A, Wodarz D, Komarova NL (2022) Aspirin’s effect on kinetic parameters of cells contributes to its role in reducing incidence of advanced colorectal adenomas, shown by a multiscale computational study. eLife 11:e71953. 10.7554/eLife.71953

Zhang R, Ukogu OA, Bozic I (2023) Waiting times in a branching process model of colorectal cancer initiation. Theor. Popul. Biol. 151:44–63. 10.1016/j.tpb.2023.04.001

